# Genomic GC bias correction improves species abundance estimation from metagenomic data

**DOI:** 10.1101/2024.09.20.614100

**Authors:** Laurenz Holcik, Arndt von Haeseler, Florian G. Pflug

## Abstract

Metagenomic sequencing measures the species composition of microbial communities, and has revealed the crucial role of microbiomes in the etiology of a range of diseases such as colorectal cancer. Quantitative comparisons of microbial communities are, however, affected by GC-content dependent biases. Here, we present GuaCAMOLE, a computational method to detect and remove GC bias from meta-genomic sequencing data. The algorithm relies on comparisons between individual species in a single sample to estimates the sequencing efficiency at levels of GC content, and outputs unbiased species abundances. GuaCAMOLE thus works regardless of the specific amount or direction of GC-bias present in the data and does not rely on calibration experiments or multiple samples. Applying our algorithm to 3435 gut microbiomes of colorectal cancer patients from 33 individual studies reveals that the type and severity of GC bias varies considerably between studies. In many studies we observe a clear bias against GC-poor species in the abundances reported by existing methods. GuaCAMOLE successfully removes this bias and corrects the abundance of clinically relevant GC-poor species such as *F. nucleatum* (28% GC) by up to a factor of two. GuaCAMOLE thus contributes to a better quantitative understanding of microbial communities by improving the accuracy and comparability of species abundances across experimental setups.

## 1 Introduction

Metagenomic sequencing has enabled the comprehensive and quantitative analysis of taxa abundances in a wide range of microbial communities [1]. It has uncovered the importance of microbiomes in health, disease, nutrition, ecology amongst others, and has revealed a complex interplay between microbial consortia and their hosts [2–4].

Metagenomic sequencing relies on comprehensive high-throughput sequencing of all DNA in a sample to quantify the abundance of all present taxa. To prepare a sample for sequencing, the DNA is extracted, purified, fragmented, amplified, and finally outfitted with sequencing adapters. Numerous protocols have been established for these library preparation steps, each differing in methods and materials used [5–7]. After sequencing, the reads are assigned to taxa and relative read counts are used as proxies for the taxa’s abundances [8, 9]. This assumes that reasonably accurate genomes of all are available. Alternatively, unknown genomes can in principle be assembled from individual reads through a process called metagenomic assembly[10–13]. Metagenomic assembly, however has a bias profile very different from that of read assignment, so we do not consider this case further here.

While metagenomic sequencing is in principle agnostic to the specific taxa in a sample, library preparation can introduce sequence-dependent biases. [14]. In particular, the GC content (i.e. fraction of G and C bases in the sequence) has been shown to strongly affect sequencing efficiency [15]. Metagenomic sequencing is particularly affected because the genomic GC content often differs significantly between species [5]. The magnitude and even direction of this bias, however, varies between different library preparation and sequencing protocols [16]. For example, a low GC content can either increase or decrease sequencing efficiency depending on the precise protocol used. As a result, computational correction for GC bias has been challenging [17].

The species on the extreme ends of the genomic GC content range are particularly prone to biases. Amongst these species we find pathogenic taxa such as *F. nucleatum* (28% GC content, associated with colorectal cancer) and *M. pneumoniae* (25% GC content, associated with pneumonia) [18–20]. With common sequencing protocols, the abundance of these taxa will be under-estimated [5, 17], and this can affect even comparisons between samples analyzed using the same protocol [21].

Ideally, GC bias should therefore be removed on a per-sample level. Computational methods to remove GC bias have been developed for various sequencing-based methods, and have been shown to be crucial to avoid skewed results [15, 22]. These methods, however, assume reads are aligned to a reference genome. For metagenomic samples possibly containing thousands of taxa, creating such an alignment is prohibitively computationally expensive. Instead, metagenomic reads are typically assigned to taxa using k-mer based methods [8, 23–25]. This makes existing methods inapplicable and requires a novel approach to GC bias correction.

We present the GuaCAMOLE (Guanosine Cytosine Aware Metagenomic Opulence Least squares Estimation) algorithm for the efficient detection and removal of GC bias from metagenomic samples. GuaCAMOLE is an alignment-free algorithm and instead assigns reads to taxa using Kraken2 [8]. The algorithm also does not require calibration data or any a-priori assumptions about the quantitative relationship between GC content and bias (such as extremely GC-rich and GC-poor species being more prone to bias), and thus works equally well for all sequencing protocols.

Using both simulations and experimental data for mock communities [16] we show that GuaCAMOLE uncovers GC bias present in samples and improves abundance estimates over Bracken [26] and MetaPhlAn4[9]. We also demonstrate that GuaCAMOLE correctly infers protocol-specific GC-dependent sequencing efficiencies.

To show that GC bias correction can be relevant in a clinical setting, we apply GuaCAMOLE to a large number of meta-genomic stool samples of colorectal cancer patients [27–29]. Here, we observe that the type and severity of GC bias varies strongly between studies, and that accounting for GC bias significantly increases the estimated abundances of clinically relevant taxa on both ends of the GC spectrum.

## 2 Results

The GuaCAMOLE algorithm processes the raw sequencing reads of a metagenomic sample and outputs bias-correct abundances for all detected taxa. GuaCAMOLE also infers and outputs GC-dependent sequencing efficiencies which reflect the probability (relative to the maximum) that a DNA fragment with a certain GC content successfully undergoes all library preparation steps and sequencing. These GC-dependent sequencing efficiencies thus measure the extent of the GC bias present in the raw data. Briefly, GuaCAMOLE works as follows (Fig. 1, see Methods for details): Reads are first assigned to individual taxa using Kraken2 [8], and within each taxa to discrete bins representing the read’s GC content (these bins are subsequently referred to as taxon-GC bins). Reads which cannot be assigned to a specific taxon unambiguously by Kraken2 are redistributed probabilistically to the likeliest taxon using the Bracken algorithm [26]. Read counts in each taxon-GC-bin are then normalized based on expected read counts computed from the genome lengths and genomic GC content distributions of individual taxa. The resulting quotients only depend on the unknown abundances (one for each taxon) and unknown GC-dependent sequencing efficiency (one per GC-bin). From these quotients we then compute bias-corrected abundance estimates and the GC-dependent sequencing efficiencies. GuaCAMOLE reports the estimated abundances either as *sequence abundances* proportional to the total amount of DNA present, or *taxonomic abundance* proportional to the number of genomes [30].

**Fig. 1.**
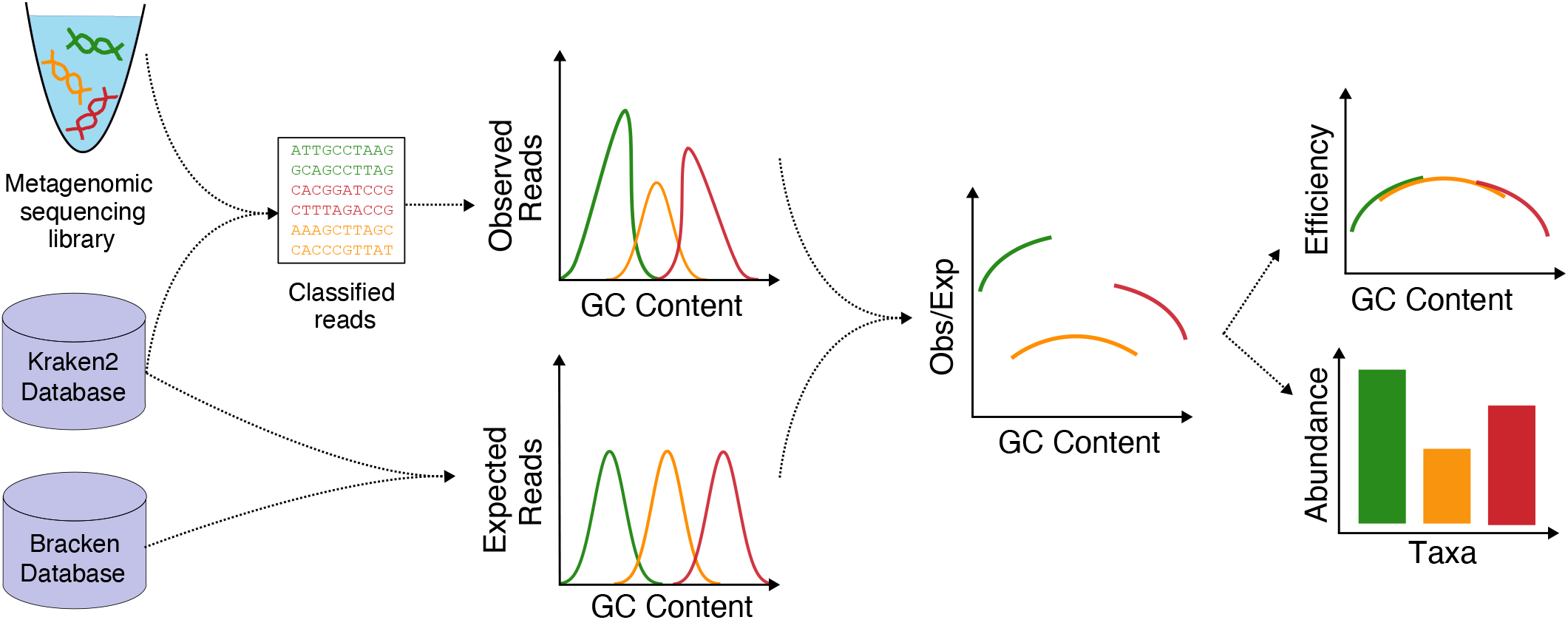
The GuaCAMOLE algorithm. Reads are assigned to taxa using Kraken2/Bracken [8, 26] and binned into discrete GC bins per taxon. Corresponding expected read counts are then computed for each taxon and GC bin from the reference genomes. The observed/expected quotients reflect the GC-dependent sequencing efficiencies scaled by each taxon’s abundance. Abundances are estimated by finding the scaling factors for which the quotients form a continuous curve.

### 2.1 GuaCAMOLE improves accuracy on simulated communities

We first demonstrate that GuaCAMOLE infers the correct abundances and GC-dependent sequencing efficiencies independent of the specific type of GC bias present. We ran GuaCAMOLE on data simulated using three different models of GC bias (see Methods for details): peak efficiency at 50% GC, efficiency increasing with GC content, and efficiency decreasing with GC content (Fig. 2A). For all three simulated datasets, GuaCAMOLE produced virtually unbiased estimates (mean relative error less than 1%) and correctly recovered the GC-dependent sequencing efficiencies used for the simulation. The Bracken estimates showed considerable GC bias in comparison (relative errors 10% to 30% depending on the bias model).

**Fig. 2.**
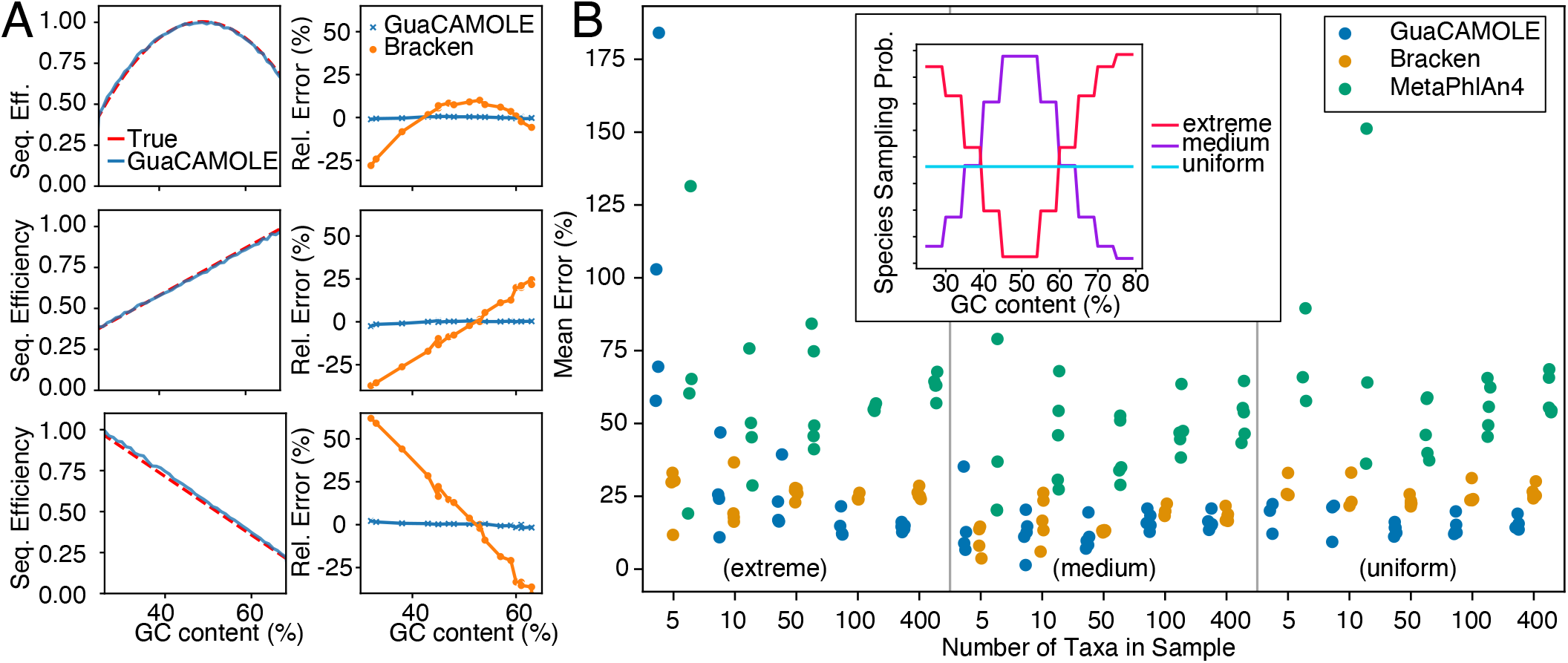
Performance on simulated metagenomic data. **(A)** Three qualitatively different types of GC-dependent sequencing bias (left column; true bias in red and GuaCAMOLE estimate in blue) and the corresponding abundance estimation errors (right column) of GuaCAMOLE (blue) and Bracken (orange). **(B)** Mean relative estimation error of GuaCAMOLE, Bracken and MetaPhlAn4 for 5 simulated communities for each combination of size (5, 10, 50, 100 and 400 taxa) and genomic GC content distributions across taxa (extreme, medium or uniform; see inset). Replicates for which GuaCAMOLE failed to produce an estimate are omitted. To avoid overlaps, dots are horizontally shifted (GuaCAMOLE left, MetaPhlAn4 right) and jittered.

We next confirmed that GuaCAMOLE performs well for metagenomic communities with different complexities and species compositions (Fig. 2B). We simulated sequencing libraries representing communities comprising different number of taxa (5, 10, 50, 100, or 400 taxa) with log-normally distributed abundances. Taxa where chosen from the RefSeq database, and selected to either have predominantly low or high genomic GC content (extreme), predominantly GC content around 50% (medium), or a GC content uniformly distributed over the whole range (uniform). For each combination of community complexity and distribution of GC content, we simulated 5 libraries with a GC-dependent sequencing efficiency of 1 – 10 · (*g −* 0.5)^2^ so that the efficiency of *g* = 30% and *g* = 70% GC content was 40%. We then compared the relative estimation errors of GuaCAMOLE with those of the other RefSeq-based algorithms, Bracken and MetaPhlAn4.

For libraries comprising 50 taxa or more, we find that GuaCAMOLE consistently shows the lowest mean estimation error (Fig. 2B). As expected, the advantage over other algorithms is the largest for communities which predominantly contain taxa with extreme GC content (extreme). When species mostly have a GC content around 50% (medium) Bracken and GuaCAMOLE perform similarly. For small communities comprising mostly taxa with extremely high or low GC content, the accuracy of GuaCAMOLE is severely reduced. For 8 out of 75 simulated communities (4 comprising 5 taxa, 3 comprising 10 taxa, one comprising 50 taxa), GuaCAMOLE was unable to produce reliable estimates and exited with a warning. The likely cause is that for these communities, the GC content distributions of individual taxa did not overlap sufficiently. Since this occurs mainly when all taxa have extreme GC content, we expect this case to be rare in practice.

### 2.2 Improved accuracy across a range of experimental protocols

Having tested GuaCAMOLE on simulated data, we went on to show that it improves abundances estimates for experimental data produced using different protocols, and that GuaCAMOLE can uncover the GC-dependent sequencing efficiencies of these protocols (Fig. 3). We re-analzyed published data [16] of a mock community sequenced using 28 different protocols (Table 1) with GuaCAMOLE, Bracken [26], MetaPhlAn4 [9], SingleM [31], Sylph [32] and mOTUS [33]. The mock community comprises 19 bacterial species representative of human-associated microbiota and was sequenced using 11 different commercially available library preparation kits (labelled A-K below, see Table 1). For each kit, Tourlousse et al. tested up to three PCR amplification regimes: 500 ng input DNA with no PCR amplification (suffix 0), 50 ng input DNA with 4-8 PCR cycles (suffix L), and 1 ng input DNA with 8-15 PCR cycles (suffix H).

**Table 1.** DNA Library Preparation Kits used for the DNA mock community [16]. Suffices 0/L/H represent no PCR amplification (0), low number of PCR cycles (L; 50 ng input DNA) and high number of PCR cycles (H; 1 ng input DNA).

| kit name | frag.<br>method | PCR<br>cycles<br>(0/L/H) | abbr. |
| --- | --- | --- | --- |
| Accel NGS 2S Plus DNA Library Kit | physical | 0 / 4 / 9 | A 0/L/H |
| QIAseq FX DNA Library Kit | enzym. | 0 / 8 / 12 | B 0/L/H |
| TruSeq (Nano) DNA (PCR-Free) Library Kit | physical | 0 / 8 / 8 | C 0/L/H |
| KAPA HTP Library Preparation Kit | physical | 0 / 5 / 15 | D 0/L/H |
| KAPA HyperPrep (PCR-free) Kit | physical | 0 / 4 / 14 | E 0/L/H |
| KAPA HyperPrep + KAPA Frag | enzym. | 0 / 4 / 14 | F 0/L/H |
| KAPA HTP + KAPA Frag | enzym. | - / 5 / 15 | G L/H |
| NEBNext Ultra II DNA Library Prep Kit | physical | - / 4 / 9 | H L/H |
| NEBNext Ultra II FS DNA Library Prep Kit | enzym. | - / 4 / 9 | I L/H |
| Nextera DNA Flex Library Prep Kit | enzym. | - / 4 / 12 | J L/H |
| SMARTer ThruPLEX DNA-Seq Kit | physical | - / 6 / 11 | K L/H |

**Fig. 3.**
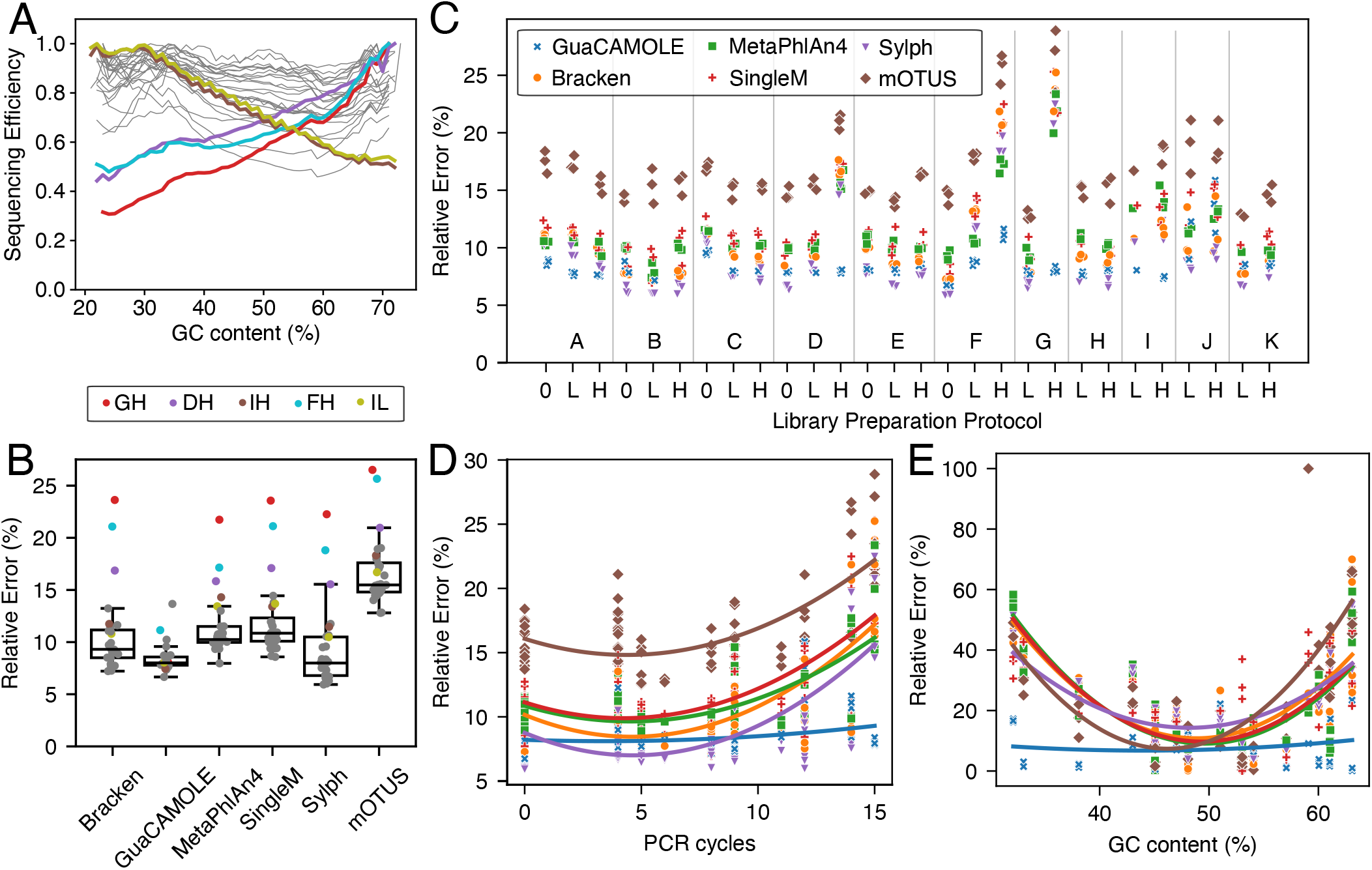
Performance of GuaCAMOLE for experimental mock community data. The mock community [16] contains 19 species and was sequenced in triplicates using 28 different library preparation protocols (Table 1). We re-analzyed the reads with GuaCAMOLE, Bracken, MetaPhlAn4, SingleM, Sylph and MOTUS. GuaCAMOLE was set to report taxonomic abundances, Bracken results were manually adjusted for genome length. Relative estimation errors are ^1^ _*j*_ |*a*_*j*_ *− A*_*j*_|*/A*_*j*_ where *A*_*j*_ is the true and *a*_*j*_ the estimated abundance of species *j* = 1, …, 19. **(A)** Estimated GC-dependent sequencing efficiencies of the 28 protocols by GuaCAMOLE. Highlighted protocols GH, DH, IH, FH, IL were found by Tourlousse et al. to exhibit the strongest dependency of efficiency on GC content. **(B)** Relative estimation errors per protocol, averaged over all replicates. Protocols GH, DH, IH, FH, IL are highlighted, see (A). Each boxplot represents 19 datapoints (one per protocol), and shows the median (center line), 25% and 75% quantiles (hinges) and the furthest point less than 1.5 IQRs (inter-quartile ranges) from the nearest hinge. **(C)** Relative estimation errors for the three replicates of each protocol. **(D)** Relative estimation error vs. number of PCR cycles used in each protocol. Lines show quadratic best fit, colors indicate the algorithm as in (C). **(E)** Relative abundance estimation error of each taxon averaged across protocols vs. genomic GC content. Lines show quadratic best fit, colors indicate the algorithm as in (C).

We find that the GC-dependent sequencing efficiencies estimated by GuaCAMOLE differ strongly between different protocols (Fig. 3A). Some protocols show uniform efficiencies, while others show a strong dependence on the GC content. In accordance with the results of Tourlousse et al. [16] we see that the protocols DH, FH, GH, IL and IH (see Table 1) show the strongest dependency on GC content (Fig. 3A, colored lines). The nature of this dependence differs qualitatively between protocols. While protocols IL and IH show decreasing sequencing efficiency with increasing GC content, DH, FH and GH show an increase in efficiency for higher GC content.

For the protocols most strongly affected by GC content (DH, FH, GH, IH, IL) GuaCAMOLE reduces the mean relative abundance error drastically compared to the other algorithms (Fig. 3B, colored dots). For other protocols GuaCAMOLE and Sylph overall show the smallest error, with a considerably larger variation of errors for Sylph than for GuaCAMOLE (Fig. 3B). Looking at individual protocols, GuaCAMOLE and Sylph show the smallest estimation errors for all protocols except protocol JH (Fig. 3C). A common source of GC bias is PCR amplification [34], and accordingly the advantage of GuaCAMOLE over the other algorithms increases with the number of PCR cycles (Fig. 3D). However, GuaCAMOLE also offers a clear advantage over the other algorithms for PCR-free protocols A0 and C0, (Fig. 3C).

The quantification error per bacterial species shows for GuaCAMOLE only a weak residual dependence on genomic GC content (Fig. 3E). In comparison, the quantification error of the other tested algorithms increase significantly for taxa on the extremal ends of the GC content range. The runtime of GuaCAMOLE (including the runtime of Kraken2 intself) is slightly longer than most other algorithms (Table S2), but the difference of up to 2x makes GuaCAMOLE still a pratical choice.

The observations that GC-dependent sequencing efficiencies are strongly affected by the choice of protocol is also observed in other datasets. For a human gut mock community [35] comprising 18 taxa sequenced on two different sequencing platforms, GuaCAMOLE reveals that the choice of sequencing platform (Illumina HiSeq 2500 rapid and NovaSeq 6000 SP) has a significant influence (Supplementary Fig. S5). Single-species libraries containing only *Fusobacterium sp. C1* [17] yields similar results (Supplementary Fig. S6).

### 2.3 GC-dependent sequencing efficiency differs widely between studies

To test how much GC-dependent sequencing efficiencies affect real-world studies, we ran GuaCAMOLE on 3435 samples from 33 studies of human gut microbiomes of healthy patients and patients with colorectal cancer (CRC), selected according to sample quality [28, 29]. 3031 of those samples were usable and reported reliable sequencing efficiencies (less than 50% reads assigned to taxa classified as false positives). By clustering these 33 studies according to the first three principal components of their average GC-dependent sequencing efficiencies (Figs. S7 and S8), we were able to identify 4 qualitatively different shapes of GC-dependent efficiency curves (Fig. 4A). With an efficiency between 50% and 100% throughout the whole GC spectrum, clusters I (10 studies) and II (16 studies) show a noticeable but limited effect of GC-content on sequencing efficiencies. In comparison, the studies in clusters III (3 studies) and IV (4 studies) exhibit a much stronger effect. Whereas in cluster III the sequencing efficiency is reduced only for GC-poor species, both GC-poor and GC-rich species are affected in cluster IV. Overall, we observe that GC-dependent sequencing efficiencies can vary drastically between otherwise similarly designed studies.

**Fig. 4.**
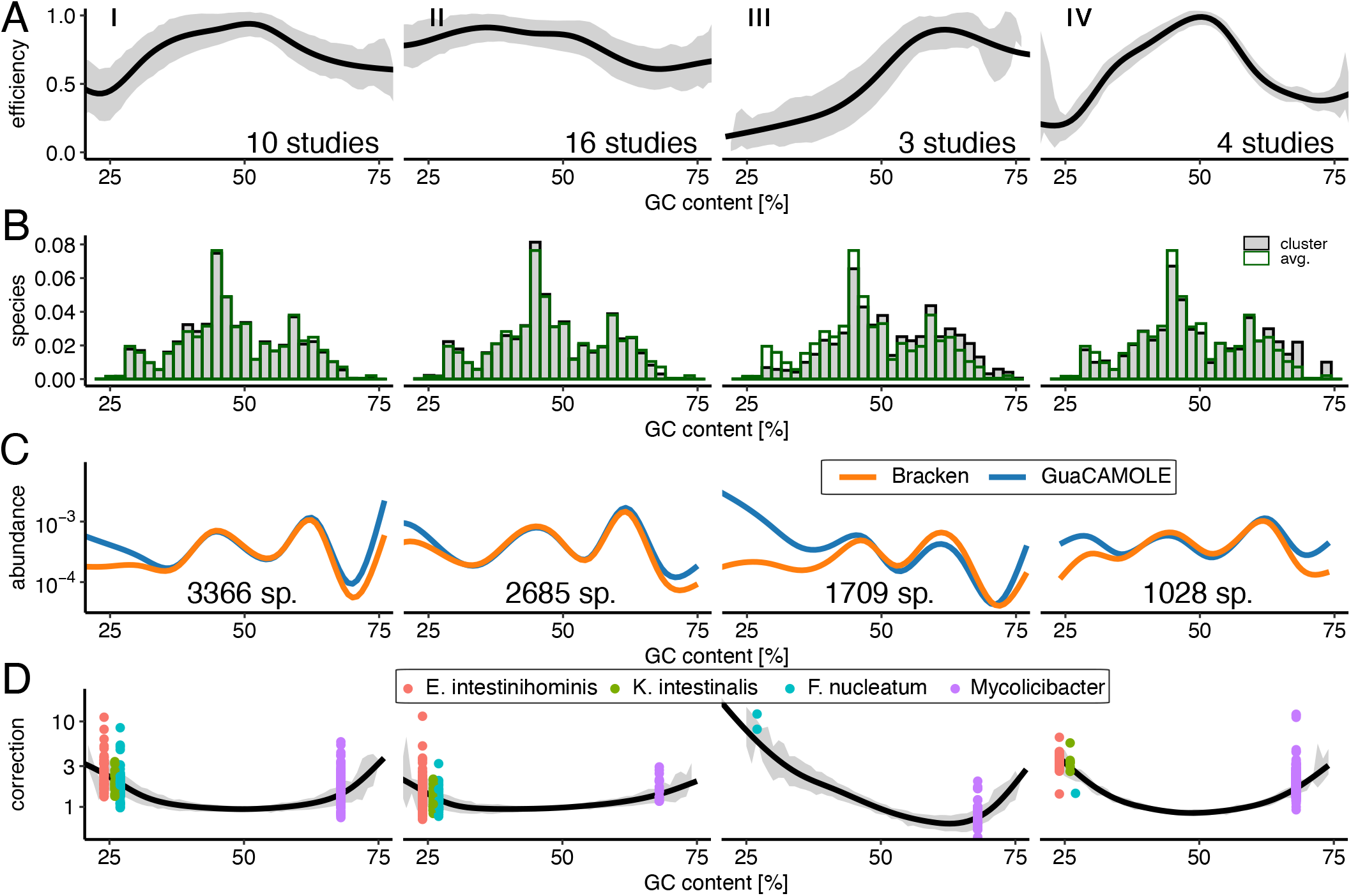
Performance of GuaCAMOLE for human gut microbiomes. GuaCAMOLE results for 3031 of 3506 samples from 34 studies of human gut microbiomes previously selected based on sample quality [28, 29]. For the remaining 475 samples GuaCAMOLE was unable to reliably estimate efficiencies. Studies were clustered into 4 clusters based on the first 3 principal components of their average GC-dependent sequencing efficiency. In panels A, C, D lines are averages across the samples in a cluster smoothed with with ggplot’s <monospace>stat_smooth</monospace>, method “GAM”. The grey areas in panels A, D ± represent one standard deviation around the average. **(A)** GC-dependent sequencing efficiencies for each of the 4 clusters. **(B)** Distribution of genomic GC content of species detected in each cluster of studies. **(C)** Average species abundance as a function of GC content. **(D)** Abundance correction applied by GuaCAMOLE.

### 2.4 Correct abundance estimation of GC-poor and GC-rich species

The distribution of average genomic GC content across detected species agrees between clusters I, II and IV, but is noticeably shifted towards higher GC content for cluster III (Fig. 4B). This indicates that the loss of sequencing efficiency for GC-poor species for studies in cluster III is severe enough to cause species to be overlooked. For GC-rich species, the effect is reversed. Here, cluster III exhibits the highest sequencing efficiency and as a result more species with high GC content are detected in cluster III than in other clusters.

The abundances of species with differing genomic GC content show a similar trend (Fig. 4C and Fig. 4D). For GC-poor speices in cluster III, the uncorrected estimates underestimate the abundance up to 10x. In clusters I and IV, both GC-rich and GC-poor species are under-estimated up to 3x.

The taxa particularly strongly affected by GC-bias are *Endlipuvirus intestinihominis* (NCBI:txid2955861), *Kahucivirus intestinalis* (NCBI:txid2956048), *Fusobacterium nucleatum* (NCBI:txid851) and *Mycolicibacter* (NCBI:txid1073531). *F. nucleatum* in particular has been associated with colorectal cancer [18–20], and is consistently underestimated by Bracken due to GC-bias; on average 1.9x in cluster I (96 samples), 1.2x in cluster II (81 samples), 10x in cluster III (2 samples) and 1.4x in cluster IV (1 sample).

GC-bias affects not only the abundance estimates of individual species, but also summary statistics about the composition of microbial communities. In particular, we observe that for most samples the alpha diversity computed from uncorrected samples is noticeably lower than after GC-bias correction (Fig. S9).

### 2.5 False positive removal

False positive taxa are created by reads which are wrongfully assigned by Kraken to species not present in the sample. Filtering detected taxa based on read counts is a common way to remove some of these false positives, but this is not always effective [36]. GuaCAMOLE additionally filters taxa based on how well their observed reads match their reference genomes by comparing observed and expected GC distributions for each taxon (see Methods). Briefly, if observed and expected read counts across a taxon’s GC bins vary more than a pre-defined threshold, the taxon is removed as an outlier, all abundances are recomputed and another round of outlier detection is initiated (see Fig. S10 for an example). This process is repeated a user-defined number of times, by default 5. This default was set based on the mock community data of Tourlousse *et al*. where it reduces the number of false-positive taxa from 18 ± 9 to 8 ± 6 and only removes a true positive in two cases (protocols JH and B0). By adjusting this thresholds, users can trade sensitivity (i.e. detecting as many taxa as possible) against specificity (avoid false positives) and accuracy of efficiency estimates (preventing false-positive taxa from interfering with the sequencing efficiency estimates).

For the simulated mock communities (Fig. 2B), GuaCAMOLE’s outlier removal reduces the number of false-positive taxa from 342 ± 218 to 120 ± 86 (Fig. S11). All together, these taxa typically represent only a few percent of all reads (Fig. S12)). The more stringent filtering of GuaCAMOLE increases the false negatives (taxa which are present but not detected) from 6 ± 10 for Bracken to 13 ± 17 for GuaCAMOLE. MetaPhlAn4 is generally much more specific and less sensitive; it finds 35 ± 54 falsepositive taxa but misses 54 ± 68 taxa actually present.

For users desiring maximal sensitivity and accuracy of abundance estimates without the risk of interference by false-positive taxa, GuaCAMOLE offers a mode which computes GC-corrected abundance estimates for all taxa detected by Bracken. In this mode, outlier removal affects only the set of taxa used to estimate GC-dependent efficiencies, and the estimated efficiencies are then used to correct the Bracken abundance estimates for GC-bias (thus effictively borrowing information between species with similar GC content). This mode was used for the analysis of the CRC samples presented in Fig. 4.

## 3 Discussion

GuaCAMOLE infers both bias-corrected abundances and GC-dependent sequencing efficiencies from a single sample without prior information about the amount or direction of GC-bias present in the data. The algorithm is agnostic about the specific sequencing protocol used and can correctly detect and correct for GC-bias without calibration data or prior knowledge about the expected type of bias. For most sequencing protocols, the bias-corrected abundances reported by GuaCAMOLE are more accurate than those reported by both Bracken and MetaPhlAn4. The advantage provided by GuaCAMOLE increases with the amount of GC bias present, and thus in particular with the amount of PCR amplification done prior to sequencing. Interestingly, we don’t observe an advantage of MetaPhlAn4 over Bracken even though we might expect the marker gene based approach of MetaPhlAn4 to be less susceptible to bias. In fact, Bracken and MetaPhlan4 often show a relatively similar quantification error.

This further corroborates that the improvement offered by GuaCAMOLE does indeed stem from successful correction of GC bias and not from other algorithmic differences.

In addition to bias-corrected abundances, GuaCAMOLE reports accurate GC-dependent sequencing efficiencies. This is useful as a quality control to check that library preparation and sequencing perform as expected. It also provides a way to estimate the amount of bias that affects taxa which remained unobserved. Finally, it allows different library preparation and sequencing protocols to be compared without the need for mock communities with known abundances.

GC bias can affect the abundance estimates of clinically relevant pathogens such as *F. nucleatum* which has been associated with a range of diseases [18–20]. We observe this in published microbiomes of colorectal cancer patients where depending on the study the abundance of *F. nucleatum* is often underestimated 2x, and can be underestimated up to 10 fold before GC bias correction. Generally we observe that the under-estimation of GC-poor and GC-rich species can differ widely between different studies of human gut microbiomes. One cause for GC-bias is likely the commonly used Nextera XT library preparation kit [5, 17]. However, the four qualitatively different shapes of GC-bias we observe in real-world human gut microbiome data suggests that many other common library preparation techniques introduce biases as well.

Even uniform biases can skew comparisons between different samples under some circumstances [21, 37]. The large qualitative difference between the GC-biases affecting different real-world human gut microbiome studies thus pose a risk for meta-analyses such as Refs. [28, 29]. While the quantitative effect of such will depend heavily on the design of such meta-analyses, GuaCAMOLE can help to mitigate these risks by uncovering the GC-bias affecting different studies, and by offering a way to correct it.

GuaCAMOLE detects false-positive taxa by checking for outliers within the deviations of observed from expected read counts. This offers more power to detect false-positive taxa than read-count thresholding, and ensures that such outliers do the skew the estimated sequencing efficiencies and abundances. However, this outlier detection assumes reasonably accurate reference genomes. Therefore, taxa which are present but whose reference genomes are inaccurate are at risk of being flagged an outlier and removed. If this is a concern, the GC-dependent efficiencies reported by GuaCAMOLE can be used to correct the bias present in the abundances estimated by other tools such as Bracken or MetaPhlAn4. This effectively “borrows” information about GC bias between species with similar GC content. For Bracken, GuaCAMOLE already implements this mode of operation as an option.

GuaCAMOLE relies on the overlap between the GC distributions of taxa to estimate sequencing efficiencies. In small communities, particularly if most taxa have an extreme genomic GC content, insufficient overlap can reduce the accuracy of GuaCAMOLE. In such cases, the fraction of reads assigned to taxa flagged as false-positives is typically also high. As a safeguard, GuaCAMOLE thus warns the user and refuses to report estimates if that fraction exceeds 50%.

The runtime of GuaCAMOLE is mostly on par with most other tools, if slightly slower. The longer runtime is to a large degree caused by the need to re-read all sequencing reads to compute their GC content. While GuaCAMOLE is even now fast enough to be pratical, a future optimization could be to modify Kraken2 to compute each read’s GC content during taxonomic mapping. Doing so would remove the need for GuaCAMOLE to access the sequencing data, and would considerably improve performance.

GuaCAMOLE provivdes a computational method to detect and correct for GC bias in sequencing protocols. For a wide range of sequencing protocols, Gua-CAMOLE substationally improves abundance estimates over alternative methods. Species whose abundance estimates are improved include clinically relevant taxa. GuaCAMOLE is in principle applicable to all types of meta-genomic samples, but performs best when reasonably complete reference genomes are available for all species. To facilitate its use by the community and its integration into standard pipelines, GuaCAMOLE is available as an easy-to-use and fast Python package under https://github.com/Cibiv/GuaCAMOLE.

## 4 Methods

### 4.1 The GuaCAMOLE algorithm

GuaCAMOLE operates on a pre-defined taxonomy comprising nodes *K*_1_, …, *K*_*n*_ which are arranged in a tree. We often refer to these nodes simply as taxa. Leaf nodes represent individual species (or strains) whereas internal nodes represent higher taxonomic groups such as genera, families etc. Leaf nodes *K*_*j*_ always have an associated genome *G*_*j*_, for internal nodes this is optional. The taxonomy together with all associated genomes is refereed to as a database.

GuaCAMOLE estimates the abundances of these taxa from a meta-genomic sequencing library containing a number *N* of sequencing reads. Each read is assumed to stem from one of the taxa in the taxonomy. The GC content of a read is the fraction of bases which are either guanin (G) or cytosin (C). We assign reads into one of *b* equally spaced bins according to their GC content, and denote the GC content by the index *g* of the respective bin.

GuaCAMOLE assumes that the composition of the sequencing library depends on (i) the abundances *a*_1_, …, *a*_*n*_ of all taxa (zero for all taxa not present in the sample), (ii) the GC-dependent sequencing efficiency *η*_*g*_, (iii) the genomic GC content distributions *f* (*j, g*) of the taxa (defined as a the expected fraction of reads from taxon *j* which fall into GC bin *g*, normalized such that Σ_*g*_ *f* (*j, g*) = 1; see section *the genomic GC content distributions* below), and (iv) the lengths *l*_1_, …, *l*_*n*_ of the taxa’s genomes. In terms of these quantities, GuaCAMOLE assumed that the number *O*(*j, g*) of fragments stemming from taxon *j* with GC content *g* in the library is

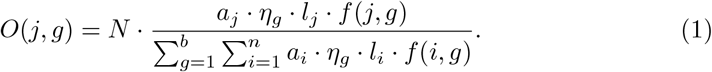

Note that abundances *a*_*i*_ and efficiencies *η*_*g*_ are defined only up to a factor by Eq. (1), we normalize these quantities by demanding that Σ_*j*_ *a*_*j*_ = max_*g*_ *η*_*g*_ = 1.

GuaCAMOLE estimates abundances and GC-dependent efficiencies by plugging observed fragment counts *O*(*j, g*), GC distributions *f* (*j, g*) and genome lengths *l*_*g*_ into Eq. (1) and solving for *a*_1_, …, *a*_*n*_ and *η*_1_, …, *η*_*b*_. Note that this system is typically over-determined: it contains on the order of *nb* equations for *n* + *b* unknowns.

### 4.2 The number of reads per taxon and GC bin

To compute observed read counts *O*(*j, g*), reads are first assigned to taxonomic nodes using Kraken2 [8]. This yields the number of assigned reads *M* (*j*) for every node *j* in the taxonomy. The reads assigned to each node are then further sub-divided according to their GC content to obtain *M* (*j, g*), the number of reads assigned to taxon *j* with GC content *g*.

The counts *M* (*j, g*), however, are biased by identical regions in the genomes of different taxa. When Kraken2 is unable to unambiguously assigned a read to a taxon due identical sequences within multiple genomes, Kraken2 assigns those reads to the lowest common ancestor (LCA) of all matching taxa. To correct for this systematic bias, we use the same approach as the Bracken algorithm introduced by Lu et al. [26]. We recall that *G*_*j*_ denotes the genome associated with node *K*_*j*_ in the taxonomy, which always exists for leaf nodes but it optional for internal nodes. Like Bracken, we compute the conditional probabilities *P* (*r ∈ G*_*j*_|*K*_*i*_) that a read which was assigned to taxon *K*_*i*_ actually stems from a descendant *K*_*j*_ of *K*_*i*_ with associated genome *G*_*j*_ in the taxonomic tree (see section 4.1). This is done by first computing *P* (*K*_*i*_|*r ∈ G*_*j*_), the probability that a read stemming from genome *G*_*j*_ is assigned to taxon *i*, by finding the taxon assigned to every possible read from genome *G*_*j*_ (of the same length as in the data). The desired probabilities *P* (*r ∈ G*_*j*_|*K*_*i*_) are then found by applying Bayes’ theorem, see Ref. [26] for details. The original Bracken algorithms uses *P* (*r ∈ G*_*j*_|*K*_*i*_) to redistribute reads assigned to *K*_*i*_. GuaCAMOLE follows the same approach, but additionally keeps track of the GC content when redistributing reads. To estimate the number of reads in GC bin *g* that stem from taxon *j*, we thus compute

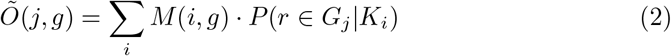

where the sum runs over all ancestors *K*_*i*_ of *K*_*j*_ in the taxonomic tree (if *K*_*i*_ is not an ancestor of *K*_*j*_, *P* (*r ∈* |*G*_*j*_ *K*_*i*_) = 0). We emphasize that the total number of reads is invariant under the redistribution done by Eq. (2); this is ensured by the fact that Σ_*j*_ *P* (*r ∈ G*_*j*_|*K*_*i*_) = 1. In particular, reads redistributed from *K*_*i*_ to a descendant *K*_*j*_ are removed from *K*_*i*_. We also note that we have assumed here for simplicity that *P* (*r∈* |*G*_*j*_ *K*_*i*_) does not depend on the read’s GC content.

To make it possible to compare abundances of higher taxonomic levels such as genera, we then sum the corrected read counts over descendants. The per-taxon and per-GC-bin read counts plugged into Eq. (1) are thus

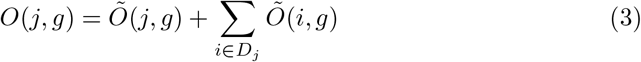

where *D*_*j*_ denotes the descendants of node *K*_*j*_.

### 4.3 The genomic GC content distributions

For Eq. (1) to hold, the observed counts *O*(*j, g*) must, in theory, arise by sampling from the the genomic GC content distributions *f* (*j, g*). These distribution must thus take the redistribution of reads into account. To find *f* (*j, g*), we first compute the individual GC distributions *q*(*j, g*) of the genomes in the taxonomy. Given the fragment length *ℓ*_*f*_ and read length *ℓ*_*r*_ of the experimental data, *q*(*j, g*) reflects the fraction of windows of length *ℓ*_*f*_ whose GC content within the parts covered by reads (i.e. the first and last *ℓ*_*r*_ bases for paired-end reads) is *g*. Here, we use the correct experimental fragment- and read length to avoid systematic errors. We then find the expected GC content distribution of the reads assigned to a specific taxon

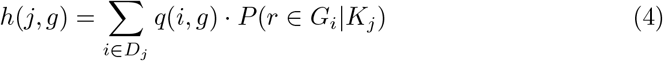

where the sum runs over the descendants of *K*_*j*_ (otherwise, *P* (*r ∈ G*_*i*_|*K*_*j*_) = 0). In Eq.4, we have thus propagated the GC distributions of individual genomes *upwards* in the taxonomic tree to account for the fact that Kraken2 will assign some reads to taxa at higher taxonomic levels. We now propagate these mixed GC distributions of internal nodes back *downwards* to find the expected GC distribution after fragment redistribution. Mimicking Eq. (2) we thus compute

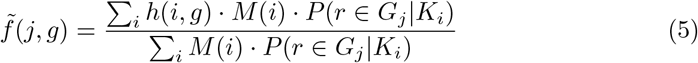

where the sums run over the ancestors of taxon *K*_*j*_. Finally, we proceed similarly to Eq. (3) and average the GC distributions of all descendants, weighted by their fragment counts,

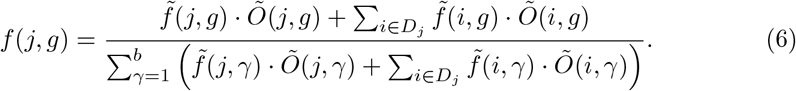

Since the tails of these distributions are typically noisy, we restrict these distributions to the range between the 2.5% and 97.5% quantile for every taxon *j*.

### 4.4 Genome lengths

We assign a single genome length *ℓ*_*j*_ to every taxon *j* independent of its taxonomic level or number of associated genomes. To do so, we average over the lengths of all assigned genomes of the taxon and all of its descendants. To account for the observed read distribution, we weight each genome length with the prior probability *P* (*K*_*j*_) that a random read stems from taxon *j* as computed by Bracken [26].

### 4.5 Estimating abundances

To estimate abundances *a*_1_, …, *a*_*n*_, we re-arrange Eq. (1) into the following expression for the GC-dependent efficiencies *η*_*g*_ in bin *g*,

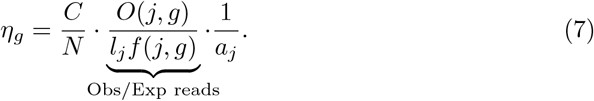

where 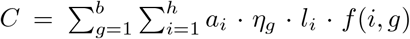 is a normalization factor. Note the correspondence to Fig. 1: For a fixed taxon *j, η*_*g*_ is proportional to the obs/exp ratio *O*(*j, g*)*/l*_*j*_*f* (*j, g*). After scaling with inverse abundances 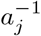 these ratios become comparable across taxa.

Given abundances *a*_1_, …, *a*_*n*_, Eq. (7) provides a separate estimate of *η*_*g*_ for every taxon whose genomic GC distribution overlaps *g*. This allows us to estimate the abundances by maximising the agreement between these separate estimates of *η*_*g*_. In terms of the inverse abundances 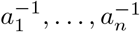 this can be expressed as minimization of the quadratic form

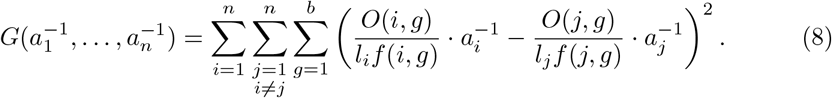

Here, we have dropped the pre-factor *C/N* from the GC-dependent efficiencies *η*_*g*_. In practice, we drop all terms from the sum in Eq. (8) that are either undefined or unreliable. A term is undefined if one of the two GC distribution does not overlap bin *g* (i.e. *f* (*i, g*) or *f* (*j, g*) is undefined). A term is considered to be unreliable if the total number of reads assigned to one of the taxa including descendants (i.e. 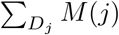 for taxon *j*, similarly for *i*) lies below some user-defined threshold (minimum read threshold, default 500).

#### Regularisation

If the taxa in a sample can be partitioned into two sets *A* and *B* such that Eq. (8) contains no term containing an abundance from *A* and a abundance from *B*, the relative abundances between sets *A* and *B* are undefined. In Fig. 1 this would be represented as two groups of taxa whose GC efficiency curves do not mutually overlap. In this situation, Eq. (8) as stated does not have a unique minimum. To avoid this, we regularize the quadratic form *G* by penalizing large differences in efficiency between neighbouring GC bins. Using Eq. (7) we express *η*_*g*_ sans the prefactor *C/N* as a weighted average of taxon-specified efficiencies,

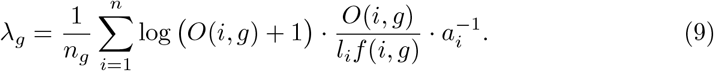

We now define the regularized objective function

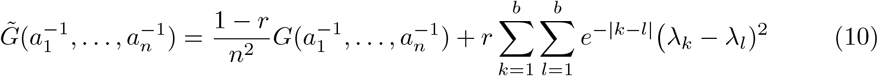

which is still quadratic since *λ*_*g*_ is linear in 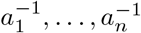. The regularized program is thus still efficiently solvable. Here, *r* is a hyperparameter that controls the amount of regularization to apply. Smaller values of *r* allow more extreme and small-scale variations in sequencing efficiency to be corrected, but increase the risk of incorrect estimates in the case of taxa partitions with non-overlapping GC distributions.

To find abundances *a*_1_, …, *a*_*n*_ we first minimizing the regularised objective function 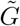 in terms of 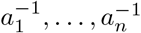 subject to 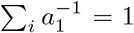 using the Python package *cvxopt*. We then compute *a*_1_, …, *a*_*n*_, normalized such that Σ_*j*_ *a*_*j*_ = 1, and compute the GC-dependent sequencing efficiencies *η*_*g*_ = *λ*_*g*_*/* max_*γ*_ *η*_*γ*_.

### 4.6 Outlier detection and removal

The set of taxa with a non-zero number *O*(*j, g*) of assigned reads often contains taxa which are not actually present in the sample. The reads witnessing such a false-positive taxon are consequently not uniformly random draws from the taxon’s genome, and we hence expect to see some deviation from Eq. (1). To detect false-positives we therefore look for outliers amongst the relative residuals *ξ*(*j, g*) = (*O*(*j, g*) *− Ō*(*j, g*)) */Ō*(*j, g*), where *Ō*(*j, g*) is the expected number of reads computed using Eq. (1). Taxa are removed if the variation *ϕ*(*j*) = max_*g*_ *ξ*(*j, g*) − min_*g*_ *ξ*(*j, g*) of *ξ*(*j, g*) of their residuals *ξ*(*j, g*) exceeds a predefined threshold *T*. After removing taxa, all abundances are recomputed, the threshold is halved, and another round of false-positive removals is done. We stop after a specified number of rounds (per default 5).

### 4.7 Simulated mock community data

We generated synthetic communities by sampling 5, 10, 50, 100 or 400 bacterial genome assemblies from RefSeq. Only assemblies of taxa which are represented with at least one genome in our Kraken2 database were considered, but we allowed the strains to differ. To control the distribution of genomic GC content within generated communities, assemblies were assigned into *N* = 12 equally sized GC-bins according to their genomic GC content and sampled in a two-stage process. First, a GC-bin *i ∈ {*1, …, 12*}* was drawn either with (i) uniform probability 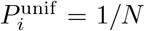, (ii) with probabilities 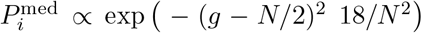 skewed towards a GC content of 50%, (iii) with probabilities 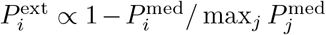 skewed towards extreme GC content. From the selected GC bin, an assembly was then drawn randomly. After sampling the genomes present in each community, the corresponding species abundances were sampled from a log-normal distribution. Finally, a paired-end sequencing library was generated for each community using a modified version of InSilicoSeq v2.0.1 [38]. In our modified version of InSilicoSeq, a bias against reads with extremal GC content is introduced by rejecting reads with probability 10 · (*g* − 0.5)^2^ where *g* is the read’s GC content.

### 4.8 Experimental mock community data

For the analysis of the mock community data of Tourlousse et al. [16] (SRA accession SRS7661134), we used the RefSeq release 220 database containing human, archaeal, viral, plasmid and bacterial DNA [39]. We ran GuaCAMOLE (in taxonomic abundance mode), Bracken and MetaPhlAn4. For Bracken and GuaCAMOLE we set the read threshold to 500. For GuacAMOLE we set the number of false-positive removal rounds to 5 (4 for JH and B0), the read length to 150bp and the fragment length to the value observed for each protocol: 200bp for D0, IL and IH, 250bp for DH, BL, BH, EH, EL, 325bp for F0, FL, FH, 350bp for JH, AH, C0, CL, 400bp for AL, A0, JL, and 300bp for all other protocols. For MetaPhlAn4 we use the CHOCOPhlAn database v202103 with default parameters, and manually corrected the classification of *F. prausnitzii* to *F. duncaniae* since this recent reclassification is not reflected by CHOCOPhlAn v202103. Since Bracken always reports sequence abundances, we adjusted the Bracken-estimated abundances using the same genome length estimates as GuaCAMOLE uses to make them comparable. For Sylph we profiled the reads using the GTDB database v220 [40]. To ensure comparability across all methods, we manually mapped the taxonomic profiles from Sylph, mOTUS3, and SingleM to their corresponding NCBI taxonomy IDs using the provided genus and species names.

### 4.9 Analyzing Colorectal Cancer Microbiome Data

We ran GuaCAMOLE on all paired-end human gut microbiom samples with non-zero read length from the curated list of samples from 33 studies found in Table S2 of Murovec *et al*. [29]. For the study of Yachida *et al*. [27] we included all 645 samples, even those not included in the list of Murovec *et al*.. GuaCAMOLE was run with default parameters except for activating genome length correction and specifying the read-length reported in the SRA meta-data of each sample. We then averaged the inferred GC-dependent sequencing efficiencies across each of the 33 studies (Fig. S7), performed a principal component analysis on the resulting efficiency curves, and clustered the studied based on the first three principal components using hierarchical clustering (R’s hclust command) with euclidean distances (Fig. S8). Based on visual inspection oft the resulting dendrogramm we split the studies into 4 clusters. For each cluster we computed the number of species, average corrected abundance, and average abundance correction compared to Bracken for each GC-bin from 25% to 75% (Fig. 4). As “corrected abundances” we used the abundances corrected based on the inferred sequencing efficiency of each taxon and we included taxa flagged as false-positives by GuaCAMOLE’s outlier detection. For plotting, abundances and abundance corrections where smoothed with R’s `stat smooth` command with method ‘GAM’.

## 5 Data availability

The SRA accessions of all samples used in this study, including the curated colorectal cancer (CRC) samples from Refs. [28, 29], together with the processed data and scripts required to reproduce the main analyses and figures of this publication are available at https://github.com/Cibiv/GenomicGCBiasCorrectionImprovesAbundanceEstimation.

## 6 Supplementary Information

**Table S2.** Runtimes for Metagenomic Analysis Tools on a Sample of 9.3 Mio read pairs. Note that Bracken as well as GuaCAMOLE require Kraken2 to be run beforehand.

| Analysis Tool | Runtime | Cores Used |
| --- | --- | --- |
| GuaCAMOLE | 4m 54s | 1 |
| Sylph | 1m 10s | 1 |
| mOTUS | 6m 25s | 10 |
| Kraken2 | 3m 27s | 10 |
| MetaPhlAn4 | 6m 11s | 10 |
| Bracken | 0.35s | 1 |

**Fig. S5.**
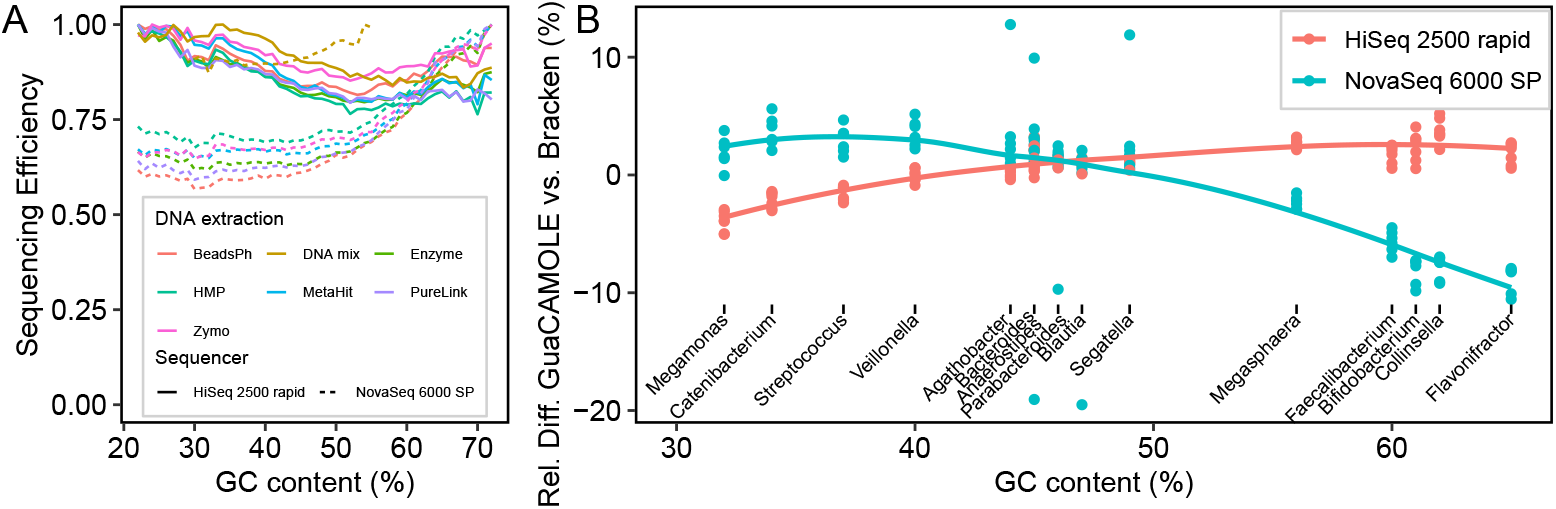
GC-bias depends on the sequencing platform. Mori *et al*. sequenced a mock community comprising 18 bacterial species with a wide range of GC contents using 7 different DNA extraction methods and two different sequencing platforms [35]. **(A)** GC-dependent sequencing efficiencies for the tested protocols and sequencing platforms. **(B)** Relative difference between the abundances reported by Bracken and GuaCAMOLE for genera with different GC content.

**Fig. S6.**
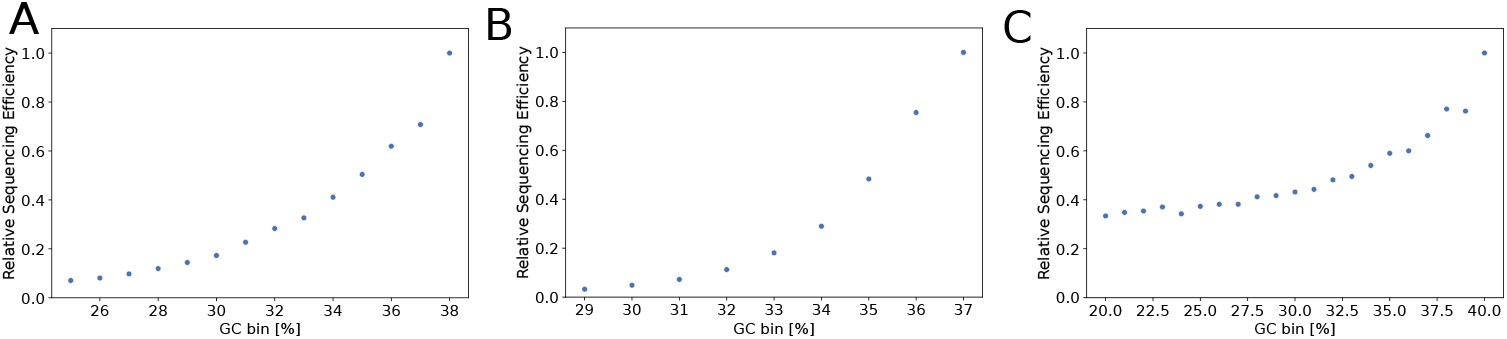
Sequencing efficiency of *Fusobacterium* in single-species libraries. Sequencing efficiencies predicted by GuaCAMOLE for single-species libraries of Browne et al. [17] containing only *Fusobacterium sp. C1* (accessions SRR8257183, SRR8257184 and SRR8257185) sequenced on different platforms. RefSeq does not contain a genome for *Fusobacterium sp. C1*, and consequently most reads were assigned by Kraken2 to *Fusobacterium hominis*. The efficiencies estimated by GuaCAMOLE agree with those reported by Browne et al. **(A)** Illumina NextSeq. **(B)** Illumina MiSeq. **(C)** Illumina HiSeq.

**Fig. S7.**
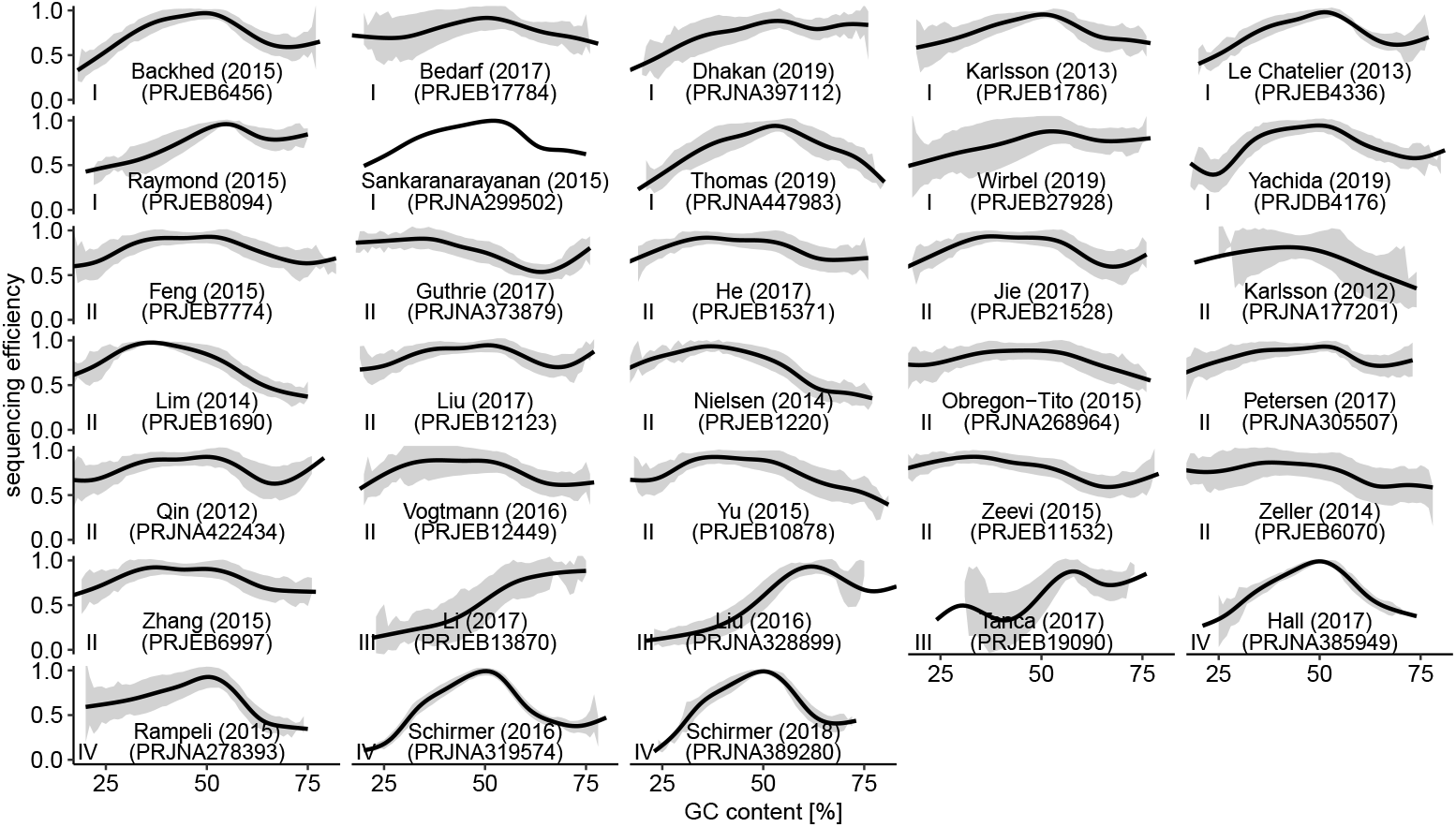
Individual GC-dependent sequencing efficiencies for the 33 studies shown in Fig. 4. Black lines are averages across the samples in each cluster smoothed with with ggplot’s stat smooth, method “GAM”. The grey areas represent *±* one standard deviation around the average.

**Fig. S8.**
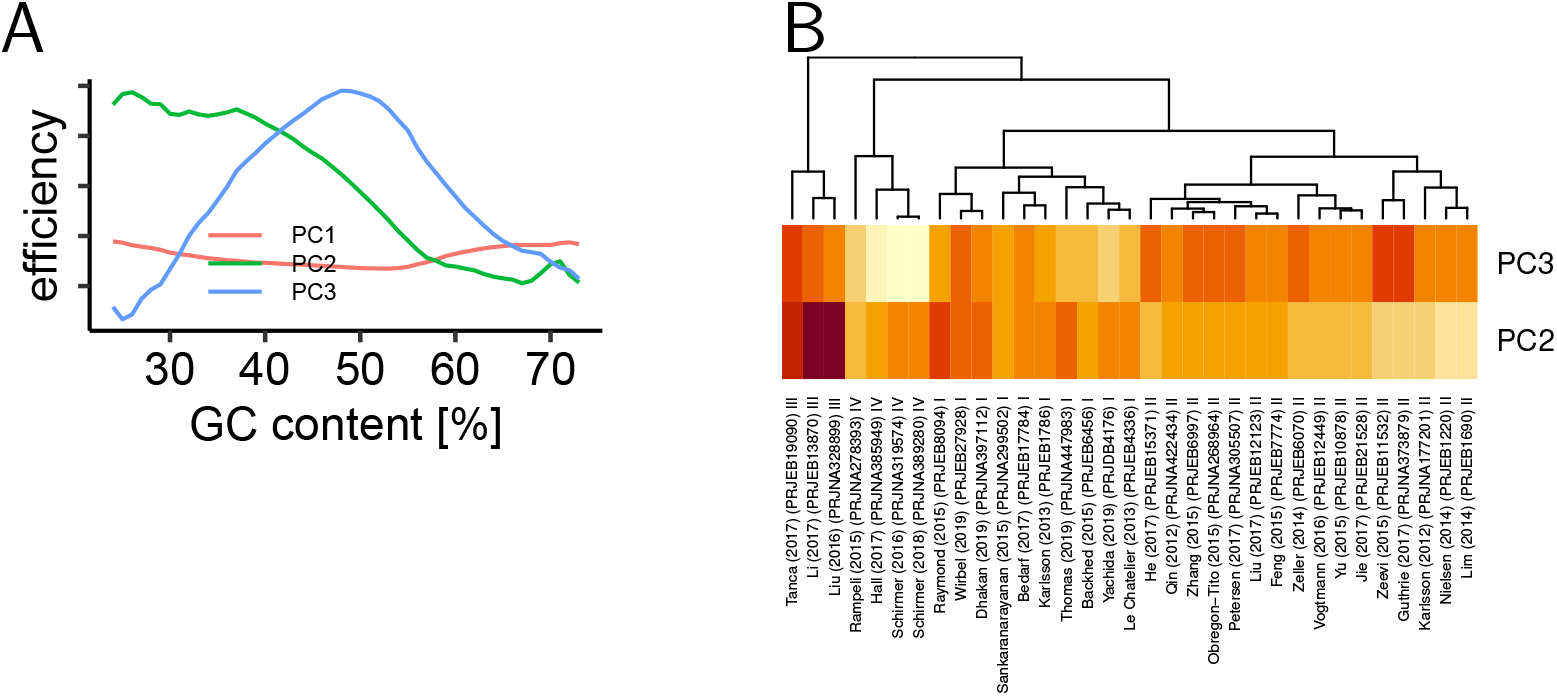
Clustering of the 33 studies shown in Fig. 4 into 4 clusters. **(A)** First three principal components of the average GC-dependent efficiencies within each study. **(B)** Hierarchical clustering based on the euclidean distances between PC1, PC2, PC3.

**Fig. S9.**
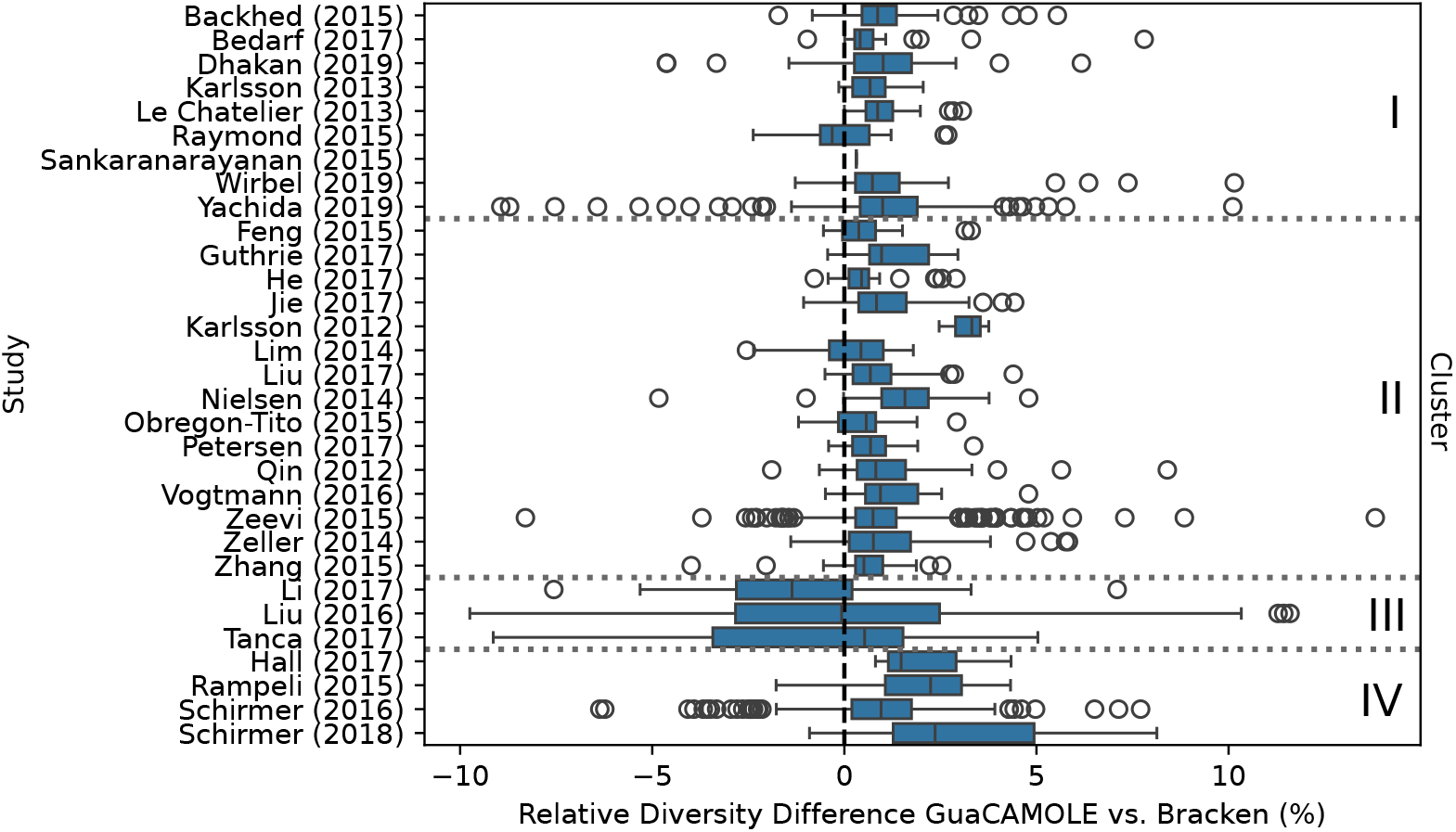
Difference in alpha diversity across studies. Alpha diversities were computed using abundance estimates by GuaCAMOLE and Bracken for each sample. Only samples of healthy individuals were used. Boxplots show the median (center line), 25% and 75% quantiles (hinges) and the furthest point less than 1.5 IQRs (inter-quartile ranges) from the nearest hinge.

**Fig. S10.**
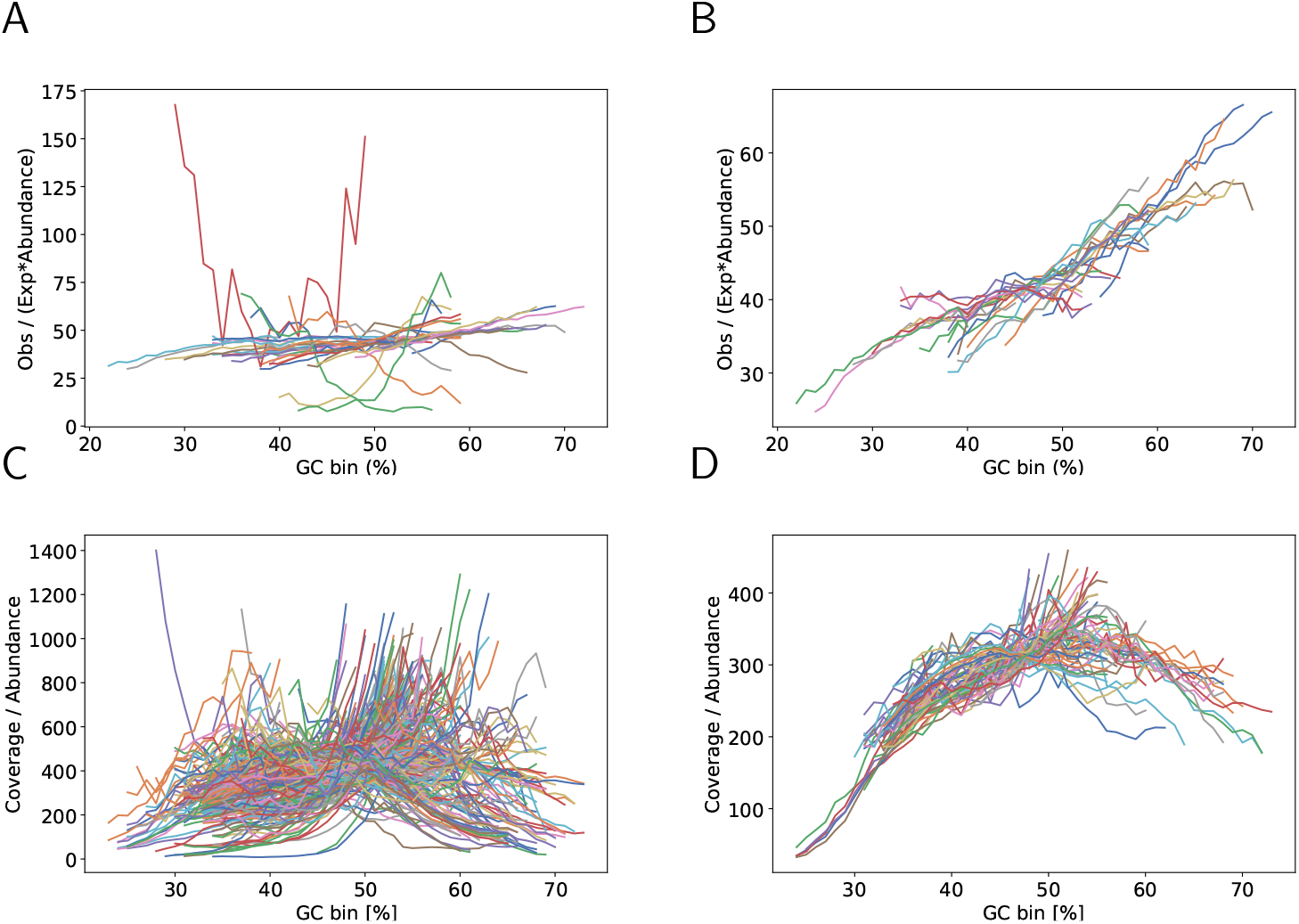
Inferred taxon-specific efficiencies before and after outlier removal. Y-axis shows *η*_*g*_ as defined in Eq. (7) without the pre-factor *C/N*. **(A)** Taxon-specific GC-dependent efficiencies for Tourlousse *et al*. protocol DH replicate b before outlier removal. **(B)** Same as (A) after 4 rounds of outlier removal. **(C)** Yachida *et al*. sample DRR127476 before outlier removal. **(D)** Same as (C) after 4 rounds of outlier removal.

**Fig. S11.**
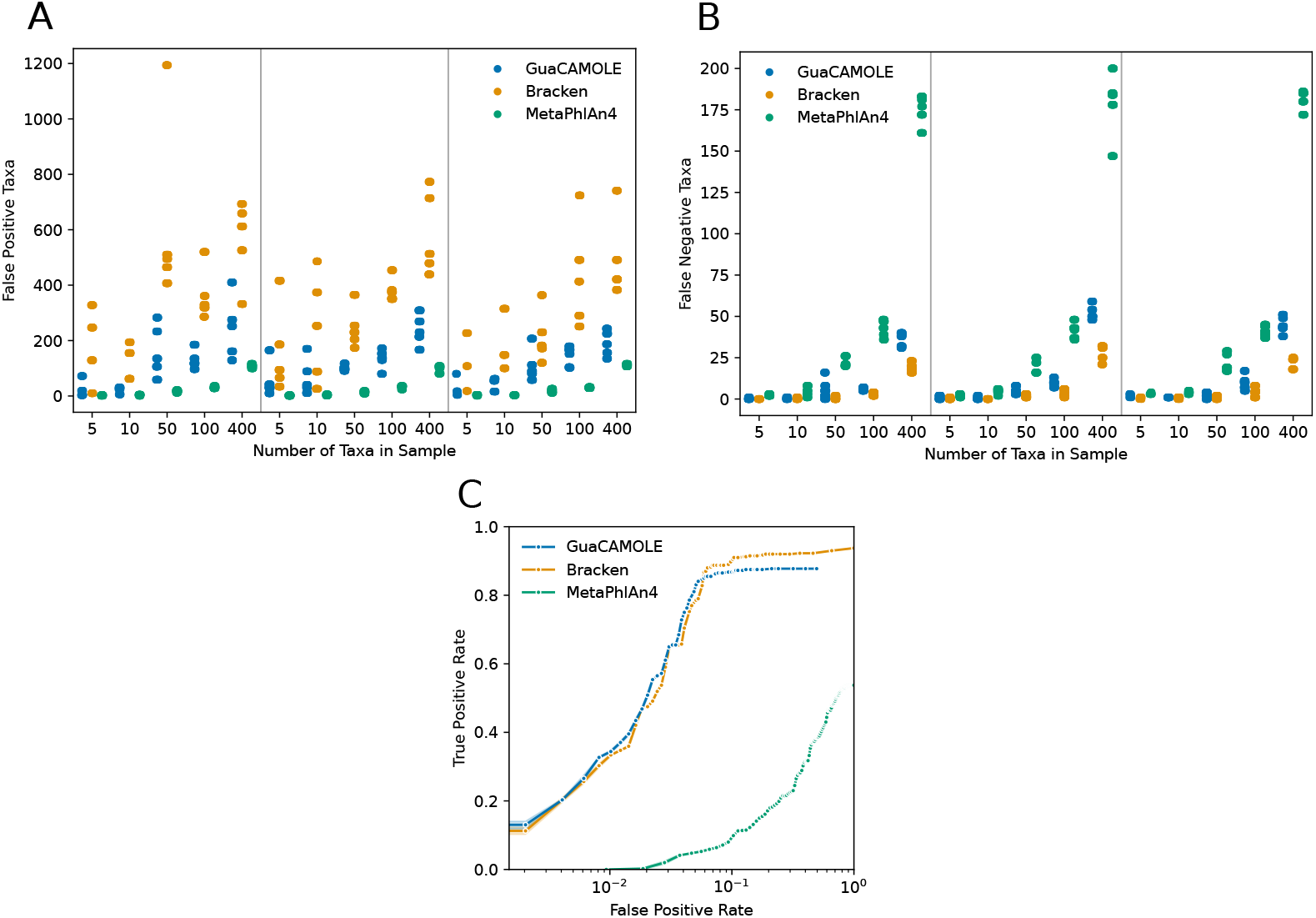
False positive and negative taxa detection of simulated communities. **(A)** Number of false positive taxa detected by GuaCAMOLE, Bracken (both with minimum read threshold threshold 100) and MetaPhlAn (with default FPKM threshold 1).

**Fig. S12.**
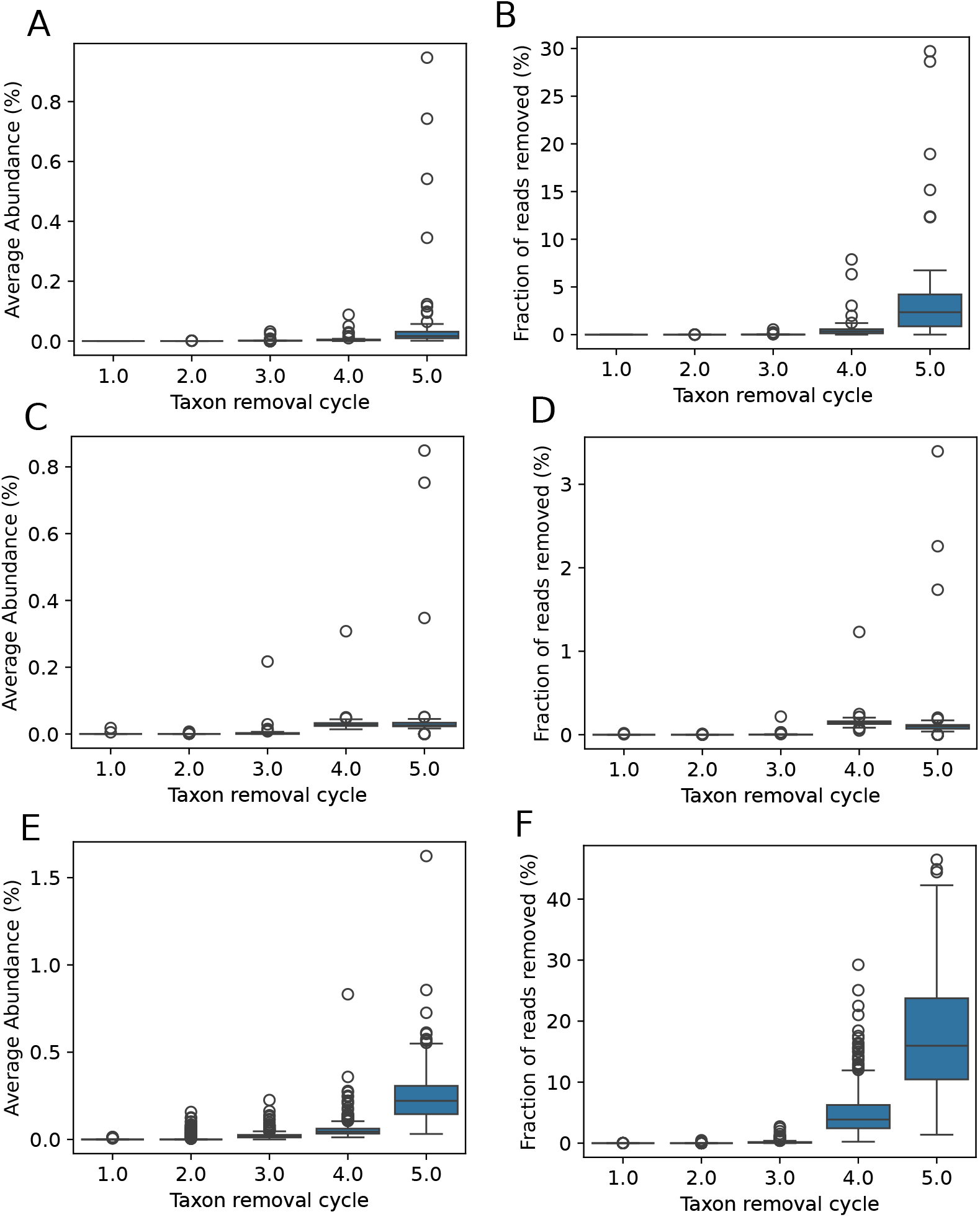
Average abundance and read fraction removed by each false positive detection cycle. Boxplots show the median (center line), 25% and 75% quantiles (hinges) and the furthest point less than 1.5 IQRs (inter-quartile ranges) from the nearest hinge. **(A)** Average abundance of taxa removed in each cycle by GuaCAMOLE for each sample of the simulated data **(B)** The fraction of total reads removed in each cycle for each cycle of the simulated data (where InSilicoSeq was used) **(C)** Same as (A) but for the [16] data set **(D)** same as (B) but for the [16]. **(E)** Same as (A) but for the [27] CRC data set **(F)** same as (B) but for the [27] CRC data set.

## References

[1] Morgan, J. L., Darling, A. E. & Eisen, J. A. Metagenomic sequencing of an in vitro-simulated microbial community. PloS one 5, e10209 (2010).

[2] Sunagawa, S. et al. Structure and function of the global ocean microbiome. Science 348, 1261359 (2015).

[3] Integrative, H. et al. The integrative human microbiome project. Natur 569, 641–648 (2019).

[4] Trivedi, P., Leach, J. E., Tringe, S. G., Sa, T. & Singh, B. K. Plant– microbiome interactions: from community assembly to plant health. Nature reviews microbiology 18, 607–621 (2020).

[5] Sato, M. P. et al. Comparison of the sequencing bias of currently available library preparation kits for illumina sequencing of bacterial genomes and metagenomes. DNA Research 26, 391–398 (2019).

[6] Bowers, R. M. et al. Impact of library preparation protocols and template quantity on the metagenomic reconstruction of a mock microbial community. Bmc Genomics 16, 1–12 (2015).

[7] Jones, M. B. et al. Library preparation methodology can influence genomic and functional predictions in human microbiome research. Proceedings of the National Academy of Sciences 112, 14024–14029 (2015).

[8] Wood, D. E., Lu, J. & Langmead, B. Improved metagenomic analysis with kraken 2. Genome biology 20, 1–13 (2019).

[9] Blanco-Míguez, A. et al. Extending and improving metagenomic taxonomic profiling with uncharacterized species using metaphlan 4. Nature Biotechnology 1–12 (2023).

[10] Ghurye, J. S., Cepeda-Espinoza, V. & Pop, M. Metagenomic Assembly: Overview, Challenges and Applications. The Yale Journal of Biology and Medicine 89, 353–362 (2016).

[11] Uritskiy, G. V., DiRuggiero, J. & Taylor, J. MetaWRAP—a flexible pipeline for genome-resolved metagenomic data analysis. Microbiome 6, 158 (2018). 10.1186/s40168-018-0541-1.

[12] MEGAHIT: An ultra-fast single-node solution for large and complex metagenomics assembly via succinct de Bruijn graph Bioinformatics Oxford Academic. https://academic.oup.com/bioinformatics/article/31/10/1674/177884.

[13] metaSPAdes: A new versatile metagenomic assembler. https://genome.cshlp.org/content/27/5/824.long.

[14] Dohm, J. C., Lottaz, C., Borodina, T. & Himmelbauer, H. Substantial biases in ultra-short read data sets from high-throughput dna sequencing. Nucleic acids research 36, e105 (2008).

[15] Benjamini, Y. & Speed, T. P. Summarizing and correcting the gc content bias in high-throughput sequencing. Nucleic acids research 40, e72–e72 (2012).

[16] Tourlousse, D. M. et al. Validation and standardization of dna extraction and library construction methods for metagenomics-based human fecal microbiome measurements. Microbiome 9, 1–19 (2021).

[17] Browne, P. D. et al. Gc bias affects genomic and metagenomic reconstructions, underrepresenting gc-poor organisms. GigaScience 9, giaa008 (2020).

[18] Yu, T. et al. Fusobacterium nucleatum promotes chemoresistance to colorectal cancer by modulating autophagy. Cell 170, 548–563 (2017).

[19] Han, Y. W. Fusobacterium nucleatum: a commensal-turned pathogen. Current Opinion in Microbiology 23, 141–147 (2015).

[20] Yang, Y. et al. Fusobacterium nucleatum increases proliferation of colorectal cancer cells and tumor development in mice by activating toll-like receptor 4 signaling to nuclear factor-κb, and up-regulating expression of MicroRNA-21. Gastroenterology 152, 851–866.e24 (2017).

[21] McLaren, M. R., Nearing, J. T., Willis, A. D., Lloyd, K. G. & Callahan, B. J. Implications of taxonomic bias for microbial differential-abundance analysis. bioRxiv 2022–08 (2022).

[22] Love, M. I., Hogenesch, J. B. & Irizarry, R. A. Modeling of rna-seq fragment sequence bias reduces systematic errors in transcript abundance estimation. Nature biotechnology 34, 1287–1291 (2016).

[23] Wood, D. E. & Salzberg, S. L. Kraken: ultrafast metagenomic sequence classification using exact alignments. Genome biology 15, 1–12 (2014).

[24] Kim, D., Song, L., Breitwieser, F. P. & Salzberg, S. L. Centrifuge: Rapid and sensitive classification of metagenomic sequences. Genome Research 26, 1721– 1729 (2016). 10.1101/gr.210641.116.

[25] Dilthey, A. T., Jain, C., Koren, S. & Phillippy, A. M. Strain-level metagenomic assignment and compositional estimation for long reads with metamaps. Nature communications 10, 3066 (2019).

[26] Lu, J., Breitwieser, F. P., Thielen, P. & Salzberg, S. L. Bracken: estimating species abundance in metagenomics data. PeerJ Computer Science 3, e104 (2017).

[27] Yachida, S. et al. Metagenomic and metabolomic analyses reveal distinct stagespecific phenotypes of the gut microbiota in colorectal cancer. Nature medicine 25, 968–976 (2019).

[28] Gupta, V. K. et al. A predictive index for health status using species-level gut microbiome profiling. Nature Communications 11, 4635 (2020).

[29] Murovec, B., Deutsch, L. & Stres, B. Predictive modeling of colorectal cancer using exhaustive analysis of microbiome information layers available from public metagenomic data. Frontiers in Microbiology 15 (2024).

[30] Sun, Z. et al. Challenges in benchmarking metagenomic profilers. Nature methods 18, 618–626 (2021).

[31] Woodcroft, B. J. et al. SingleM and Sandpiper: Robust microbial taxonomic profiles from metagenomic data (2024).

[32] Shaw, J. & Yu, Y. W. Rapid species-level metagenome profiling and containment estimation with sylph. Nature Biotechnology 1–12 (2024). 10.1038/s41587-024-02412-y.

[33] Ruscheweyh, H.-J. et al. mOTUs: Profiling Taxonomic Composition, Transcriptional Activity and Strain Populations of Microbial Communities. Current Protocols 1, e218 (2021). 10.1002/cpz1.218.

[34] Aird, D. et al. Analyzing and minimizing pcr amplification bias in illumina sequencing libraries. Genome biology 12, 1–14 (2011).

[35] Mori, H. et al. Assessment of metagenomic workflows using a newly constructed human gut microbiome mock community. DNA Research 30, dsad010 (2023). 10.1093/dnares/dsad010.

[36] Bradford, L. M., Carrillo, C. & Wong, A. Managing false positives during detection of pathogen sequences in shotgun metagenomics datasets. BMC bioinformatics 25, 372 (2024).

[37] McLaren, M. R., Willis, A. D. & Callahan, B. J. Consistent and correctable bias in metagenomic sequencing experiments. Elife 8, e46923 (2019).

[38] Gourlé, H., Karlsson-Lindsjö, O., Hayer, J. & Bongcam-Rudloff, E. Simulating illumina metagenomic data with insilicoseq. Bioinformatics 35, 521–522 (2019).

[39] O’Leary, N. A. et al. Reference sequence (refseq) database at ncbi: current status, taxonomic expansion, and functional annotation. Nucleic acids research 44, D733–D745 (2016).

[40] Parks, D. H. et al. Gtdb: an ongoing census of bacterial and archaeal diversity through a phylogenetically consistent, rank normalized and complete genome-based taxonomy. Nucleic acids research 50, D785–D794 (2022).

